# Tuning Scaffold Degradation with Non-Natural Peptidomimetics to Control Human Umbilical Vein Endothelial Cell Morphology and Vessel Formation

**DOI:** 10.1101/2025.10.27.684851

**Authors:** Kathleen N. Halwachs, Carolyn M. Watkins, Morgan J. Vadivieso, Janet Zoldan, Adrianne M. Rosales

## Abstract

The in vitro vascularization of 3D tissue constructs, such as hydrogels, remains a paramount challenge in tissue engineering. Extracellular matrix degradation and remodeling are key parts of the vascularization process; however, it is difficult to isolate the effects of degradability in both natural and synthetic matrix models. Naturally-derived matrices typically couple degradability to other material properties, whereas synthetic matrices rely on short peptide sequences to impart degradability, which typically exhibit substrate overlap to many proteases. Here, we present a method to independently and broadly tune 3D hydrogel degradation using crosslinkers with non-natural peptoid (N-substituted glycine) substitutions. Increased peptoid substitutions reduced hydrogel degradability to collagenases without altering hydrogel modulus, swelling ratio, or crosslinker length. Using this approach, human umbilical vein endothelial cells (HUVECs) encapsulated in more degradable hydrogels proliferated more, formed more vessels, exhibited higher metabolic activity, and secreted more extracellular matrix than HUVECs encapsulated in less degradable or non-degradable hydrogels. Interestingly, HUVECs encapsulated in the least degradable hydrogels secreted significantly higher matrix metalloproteinase-2 (MMP-2) and matrix metalloproteinase-9 (MMP-9) than HUVECs encapsulated in the most degradable hydrogels, suggesting higher MMP secretion to compensate for the reduced matrix degradability. Overall, this work highlights the importance of protease-mediated remodeling on vascularization and suggests that peptoid substitutions are effective for tuning hydrogel degradability for a variety of 3D cell applications.

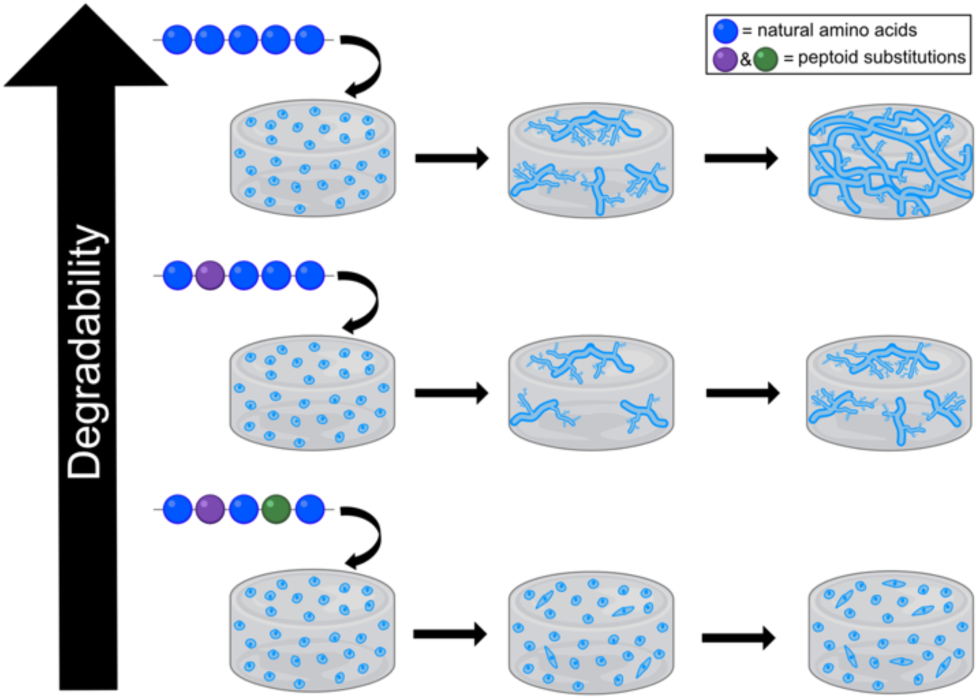

## 1. Introduction

The vascularization of tissue constructs remains a critical bottleneck of tissue engineering.^1^ Specifically, the generation of perfusable, interconnected vascular networks are vital for maintaining homeostasis and transporting oxygen, nutrients, and waste products.^2^ One approach for vascularizing tissue constructs involves encapsulating endothelial cells (ECs) within a scaffold that promotes vasculogenesis, or the de novo assembly of endothelial cells into vessels.^3^ The process of vasculogenesis requires substantial matrix remodeling because the ECs must adjust the surrounding scaffold to allow for migration, proliferation, and the assembly of new vessels.^4,5^ To support the formation of vascular networks, the scaffold must balance adaptability and mechanical integrity; it must be robust enough to support the newly forming vessels while being degradable enough to allow for network formation.

Degradability to matrix metalloproteinases (MMPs) has been well studied in many biomaterial platforms as a moderator of vascular network formation.^6,7^ Previous work leveraged naturally degradable polymers, specifically collagen, fibrin, and gelatin, for vasculogenic scaffolds by altering the polymer concentration or the amount of crosslinking, finding that more degradable matrices result in enhanced vessel formation.^8–13^ However, altering the polymer concentration or the amount of crosslinking not only impacts scaffold degradability, but it also alters the matrix stiffness, the network permeability, and the densities of cell adhesion sites.^14^ Given that these material parameters are coupled, other studies have used synthetic polymers, such as poly(ethylene glycol) (PEG), with short, MMP-degradable peptide sequences of different identities or with different numbers of repeats of the sequence.^15–21^ While this approach decouples degradability from adhesion, many short peptide sequences suffer from overlapping cleavage specificity profiles.^22,23^ Since these short peptide sequences do not degrade to specific enzymes and cells secrete a mix of proteases at unknown concentrations, changing the identity of the crosslinker can result in slower degradation to some proteases and faster degradation to others and often does not result in major differences in vessel formation.^17,21^ Notably, a recent high-throughput approach combining proteomics, degradation studies, and split-and-pool synthesis had success generating differentially-degradable short peptide crosslinkers that resulted in significant differences in vessel formation; however, this approach is very laborious.^24^ Thus, a streamlined, rational design approach to broadly decrease hydrogel degradability to proteases while keeping material properties tightly controlled is necessary.

Recently, our group developed degradable peptide-based molecules that leveraged non-natural, peptoid substitutions to systematically modulate degradability.^25–27^ Peptoids, or N-substituted glycines, reduce the proteolytic susceptibility of a peptide sequence by displacing key side chains from the α-carbon to the amide nitrogen. Increased peptoid substitutions decrease proteolytic susceptibility further across a broad range of proteases, avoiding confounding effects from sequences optimized for specific proteases.^27,28^ In addition, previous work found peptoids or peptoid-containing molecules to be cytocompatible,^29–32^ though this property is sequence-dependent. In this study, we leveraged peptoid substitutions in a degradable peptide sequence to modulate cleavage rate while keeping the overall peptide length and chemical composition fixed. We used these sequences to crosslink cell-laden hydrogels of hyaluronic acid (HA), which is a natural ECM component known to play a role in vessel formation by influencing cell migration, proliferation, and tube formation.^33–35^ HA is also easily functionalized to incorporate cell adhesive motifs.^36,37^

We encapsulated human umbilical vein endothelial cells (HUVECs) in the HA hydrogels and quantified cell viability, vessel formation, secretion of MMP-2 and MMP-9, cell morphology, and extracellular matrix (ECM) secretions. MMPs, particularly MMP-2 and MMP-9, and the secretion of nascent ECM have been established as critical in vasculogenesis.^38^ MMPs are responsible for ECM degradation which allows EC migration, branching, lumen formation, and the release of several cytokines which are known to influence vessel formation.^7,38,39^ Additionally, the ECM provides organizational cues to ECs, structural support to vessels, helps maintain vascular homeostasis, and serves as a reservoir for several cytokines involved in vessel formation.^6,40^ Thus, quantifying the proteolytic activity of MMP-2 and MMP-9 from encapsulated HUVECs and the ECM secretions of individual HUVECs provided insight into the mechanisms through which degradability was impacting vessel formation in this hydrogel system. Altogether, this study presents a hydrogel platform that independently tunes hydrogel degradation with non-natural peptidomimetic residues to a range of proteases, quantifies the significant effect of degradability on vessel formation, and investigates why differences in vessel formation are observed.

## 2. Materials and Methods

### 2.1 Norbornene functionalized hyaluronic acid (NorHA) synthesis

Norbornene functionalized hyaluronic acid (NorHA) was synthesized similarly to what has been described previously.^41^ In brief, we dissolved HA (Na-HA, 72 kDa, Lifecore Biomedical) in DI water at 2 wt% with Dowex 50W ion exchange resin (4:1 resin to HA by weight, Millipore Sigma) and stirred the mixture for 5 hours at 600 RPM. Next, we filtered out the Dowex resin, titrated the solution to pH 7.03 with tetrabutylammonium hydroxide (0.4 M, Thermo Scientific), froze the solution (-80°C), and lyophilized it. Then, we dissolved HA-TBA, 5-norbornene-2-carboxylic acid (5-NB-2-CA, Thermo Scientific) (7.5:1 M ratio of 5-NB-2-CA to HA-TBA repeat unit) and 4-(dimethylamino)pyridine (DMAP, Acros Organics) (3.75:1 M ratio of DMAP to HA-TBA repeat unit) in anhydrous dimethyl sulfoxide (DMSO) under argon at 45°C. One hour later, di-tert-butyl dicarbonate (Boc_2_O, Thermo Scientific) (1:1 M ratio of Boc_2_O to HA-TBA repeat unit) was added to the flask through a syringe, and the reaction was allowed to proceed for 24 hours. The reaction was quenched with 4x excess cold DI water (4°C) and dialyzed using a Spectra/Por membrane with a molecular weight cutoff of 6-8 kDa in DI water for three days with daily water changes. After three days, 1 g sodium chloride (Fisher BioReagents) per 100 mL was added and the solution was precipitated into 6X excess cold acetone (-20°C, Fisher Chemical). The precipitate was centrifuged, the supernatant acetone was decanted, the pellet was dried, and then the pellet was re-dissolved in DI water and dialyzed against DI water for another seven days with daily water changes. After dialysis, the solution was frozen (-80°C), lyophilized, and then stored at -20°C. Following lyophilization, ^1^H NMR (deuterium oxide) confirmed that 48% of repeat units had been functionalized with norbornenes (**Fig. S1**).

### 2.2 Peptide/peptoid synthesis and purification

RGD (GCGYGRGDSPG) and peptide/peptoid crosslinkers were all synthesized by a Prelude X automated peptide synthesizer (Gyros Protein Technologies) on Rink Amide polystyrene resin (0.48 mmol g^-1^, Chem-Impex) at a scale of 250 μmol. Fmoc groups were removed from the Rink Amide resin and subsequent amino acids by washing twice with 20% piperidine (Millipore Sigma) in dimethylformamide (DMF). Fmoc-protected amino acids (250 mM, 5.0 equiv., Chem-Impex) were coupled using O-(1H-6-chlorobenzotriazol-1-yl)-N,N,N′,N′-tetramethyluronium hexafluorophosphate (HCTU) activator (250 mM, 5.0 equiv., Chem-Impex) and N-Methylmorpholine (NMM, 500 mM, 10.0 equiv., Millipore Sigma). Coupling steps were performed twice. Upon completion of synthesis, peptides, peptomers, and peptoids were cleaved from the Rink Amide resin by mixing it for 2-4 hours with cleavage cocktail (94:2.5:2.5:1 %(v/v) trifluoroacetic acid:ethane-1,2-dithiol:water:triisopropylsilane). The resin was then filtered off, and the peptides, peptomers, and peptoids were precipitated into a 10-fold volume of chilled (-20°C) diethyl ether (Fisher Scientific), washed 2x with fresh diethyl either, then centrifuged to collect the product. Peptides/peptoids were dissolved in a mixture of acetonitrile (Fisher Scientific) and water (ranging from 10-40% acetonitrile) with 0.1% trifluoroacetic acid, filtered with a PTFE 0.45 µM filter, and purified using a semi-prep C18 column on a Dionex UltiMate 3000 UHPLC with a 15 minute gradient from starting composition to 100% acetonitrile at 10 mL min^-1^, with gradient adjustments as needed. Peptides, peptomers, and peptoids were collected by their 214 nm UV signal, frozen (-80°C), and lyophilized. To verify molecular weight, final products were analyzed with matrix-assisted laser desorption/ionization time-of-flight (MALDI-TOF) mass spectrometry using a Bruker autoflex maX instrument (**Fig. S2-S7**).

### 2.3 Rheology

*In situ* gelation mechanics were measured on a Discovery HR-2 rheometer (TA Instruments). For all experiments, an 8 mm flat stainless-steel geometry was used with a UV-transparent quartz plate. The UV transparent quartz plate was connected via light guide to a mercury lamp (Omnicure Series 1500) fitted with a 365 nm filter. The hydrogel precursor solution (20 µL) was pipetted onto the quartz plate and exposed to 365 nm light (10 mW/cm^2^, 50 s) to induce gelation. Time sweeps were performed at 1 rad/s and 1% strain. At least four hydrogels per condition had their storage modulus measured.

### 2.4 Swelling Study

To compare the swelling ratios of hydrogels made with different crosslinkers, we fabricated 50 µL hydrogels by pipetting hydrogel precursor solution onto a silicone sheet and crosslinking the hydrogels via exposure to 365 nm light (10 mW/cm^2^, 50 s) from a mercury lamp (Omnicure Series 1500) connected to a collimator. Upon gelation, the hydrogels were transferred to a 24 well plate and 1.5 mL of deionized water was added to each well. The hydrogels swelled for 48 hours at 37°C, and the swollen mass was obtained by weighing each hydrogel in a pre-massed microcentrifuge tube. Then, the hydrogels were frozen overnight in the microcentrifuge tubes and lyophilized until fully dry. The microcentrifuge tubes containing the dried hydrogels were then weighed again to determine the dry mass. At least four hydrogels per condition had their swelling ratio calculated. The dry and swollen masses were used to determine the equilibrium swelling ratio of each hydrogel using Equation 1:

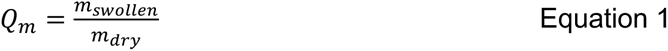

### 2.5 Collagenase Degradation Study

To quantify hydrogel degradation to collagenase, 30 µL hydrogels were fabricated on a silicone surface and transferred to a 24 well plate in which they were swollen in 1.5 mL of PBS for 48 hours at 37°C. The hydrogels were massed, and the PBS was replaced with 1 mL of collagenase solution. The collagenase solution consisted of 250 units/mL collagenase type I (isolated from Clostridium histolyticum, activity of 250 U/mg, ThermoFisher Scientific) dissolved in buffer containing 50 mM tricine, 10 mM calcium chloride, and 400 mM sodium chloride. Then, hydrogels were incubated at 37°C and removed and weighed at many timepoints to determine the remaining mass. The buffers were replaced with fresh solution every 24 hours. At least three hydrogels were massed per condition at every timepoint.

### 2.6 Hyaluronidase Degradation Study

To quantify hydrogel degradation to hyaluronidase, 50 µL hydrogels were fabricated on a silicone surface and transferred to a 24 well plate in which they were swollen in 1.5 mL of PBS for 48 hours at 37°C. The hydrogels were massed, and the PBS was replaced with 1 mL of hyaluronidase solution. The hyaluronidase solution contained 75 units/mL of hyaluronidase isolated from bovine testes (MP Biomedical’s) dissolved in 0.1 M sodium phosphate buffer with 0.15 M sodium chloride. Then, hydrogels were incubated at 37°C and removed and weighed at many different timepoints to determine the remaining mass. At least three hydrogels were massed per condition at every timepoint.

### 2.7 Cell Culture

Human umbilical vein endothelial cells (HUVECs) were purchased from PromoCell (catalog number C-12203, lot number 488Z001) at passage 1. They were cultured on T75 tissue culture plastic flasks in Endothelial Cell Growth Medium 2 (PromoCell, catalog number C-22011) supplemented with 1 v% penicillin/streptomycin (Corning). Media changes occurred every other day and cells were cultured in an incubator (at 37 °C and 5 v% CO_2_). Once the HUVECs reached approximately 80% confluency, they were lifted for five minutes at room temperature with 0.05% trypsin, 0.53 mM EDTA solution (Corning). The trypsin was neutralized with an equal volume of Corning’s Dulbecco’s Modification of Eagle’s Medium (DMEM) supplemented with 10 v% fetal bovine serum (Corning) and 1 v% penicillin/streptomycin (Corning) and then the HUVECs were pelleted out via centrifugation for 3 minutes at 220 RCF.

### 2.8 Cell Encapsulation

Cells were encapsulated between passages 3 and 5 at a density of 2.2x10^6^ cells/mL for all experiments except for metabolic labeling experiments. For metabolic labeling encapsulations, a density of 1x10^5^ cells/mL was used. NorHA was first dissolved in PBS to a final concentration of 1 wt% and then pre-dissolved RGD was added to the NorHA solution to a final concentration of 2 mM. Next, pre-dissolved photoinitiator lithium phenyl (2,4,6-trimethylbenzoyl) phosphinate (LAP) was added to a final concentration of 0.05 wt%. Crosslinker was added at a concentration of 1.36 mM to theoretically achieve 25% crosslinking of available norbornene functional groups. After passaging and centrifuging into a pellet, cells were resuspended in 10 μL of PBS and added to the precursor solution. All components were then mixed by pipetting, and pipetted so that the hydrogel formed a bead on the surface of a tissue culture plastic 24 well plate. The solution was then exposed to 10 mW/cm^2^, 365 nm light for 50 s. Immediately after gelation, EGM2 complete media supplemented with 1 v% penicillin/streptomycin and 50 ng/mL of VEGF165 (Bon Opus Biosciences) was added. Media was changed daily during 3D culture.

### 2.9 Live/Dead Assay, Fluorescent Imaging, and Quantification

Live/Dead (Invitrogen) assays were performed one-hour post-encapsulation for all crosslinking conditions. Cells were encapsulated as described in Section 2.8; after an hour of incubation, the media was removed and the hydrogels were washed with PBS 2x5 minutes. Then, the PBS wash was replaced with a PBS solution containing 2 μM calcein-AM and 4 μM ethidium homodimer-1, and the hydrogels were incubated at 37°C and 5 v% CO_2_ for 30 minutes. The cell staining solution was removed, the hydrogels were washed with fresh PBS for five minutes, and the cells were imaged with a 10X objective on a Nikon Eclipse Ti2 microscope. Cells in which the membrane-permeable calcein-AM was cleaved were counted as live, while cells permeable to ethidium homodimer-1 were counted as dead. Over 100 cells were counted per image. For each condition, three separate encapsulations were performed. One hydrogel was imaged (in three different locations) per condition from each encapsulation. To quantify percent viability, maximum intensity projections of the calcein-AM channel and the ethidium homodimer-1 channel were generated, thresholded, and binarized in ImageJ. Then, the ImageJ particle analysis command was used to count the number of cells in each channel. Percent viability was calculated by dividing the number of cells in the calcein-AM channel by the total number of cells (the number of cells in the calcein-AM channel added to the number of cells in the in the ethidium homodimer-1 channel).

### 2.10 MTS Viability Assay

A colorimetric MTS viability assay (Promega) was performed at day 3 and day 7 after encapsulating cells as described in Section 2.8. On the day that the assay was performed, hydrogels were transferred into a new 24 well plate along with 500 µL of fresh media and 100 µL of MTS reagent. The well plate was incubated for 4 hours at 37 °C and 5 v% CO_2_, then the contents of the wells were mixed well via pipetting, and the absorbance was read at 490 nm using a BioTek Synergy H1 Microplate reader.

### 2.11 Metabolic Labelling and Analysis

To analyze cell morphology and use metabolic labelling to quantify nascent ECM secretions, we followed a pre-established protocol.^42,43^ To do this, we encapsulated cells at a density of 100,000 cells/mL (as described in Section 2.8) and supplemented the media with 50 μM N-Azidoacetylmannosamine-tetraacylated (Sigma Aldrich). After three days, hydrogels were washed thrice with PBS for 5 min at 37°C. The third wash was replaced with 1% bovine serum albumin (BSA, Thermo Scientific) in PBS containing 30 μM DBCO-Cy5 (AAT Bioquest), and the hydrogels were incubated for 40 min at 37°C. Hydrogels were then washed with PBS 3x5 min at 37°C to remove any excess DBCO-Cy5 and then fixed for one hour at room temperature with 4 v% paraformaldehyde (Thermo Scientific) in PBS. Following fixing, hydrogels were washed with PBS 3x5 min at room temperature and the final wash was replaced with a permeabilization solution of 0.2 v% Triton X-100 (Millipore Sigma) in 2 v% paraformaldehyde solution. After permeabilization for two hours at room temperature, hydrogels were again washed with PBS 3x5 min at room temperature. The final wash was replaced with a blocking buffer comprised of 5 v% BSA, and the samples were incubated at room temperature for two hours. Then, the blocking buffer was replaced with a staining solution comprised of 1 v% BSA, DAPI (1:500, Sigma Aldrich), and CellBrite Orange (1:200, Biotium), and the samples were stained overnight at 4°C. The next day, all hydrogels were washed 5x5min with PBS before being imaged on a Nikon AXR-NSPARC with a 20X air objective and 6X zoom. Imaging was performed by placing the hydrogels in #1.5 glass bottom mini petri dishes (Cellvis). Three encapsulations were performed, and three hydrogels were imaged per condition from each encapsulation. Five cells were imaged in each hydrogel for a total of forty-five cell images per condition. To quantify average thickness of nascently deposited extracellular matrix (ECM), ImageJ was used to generate maximum intensity projections of the CellBrite Orange membrane channel and the DBCO-Cy5 channel.^44,45^ Then, Otsu’s thresholding was used to binarize these maximum intensity projections, and, to avoid quantification of intracellular staining, the cell membrane mask was subtracted from the DBCO-CY5 mask. Then, the analyze particles command was used to compute the total area of the nascent ECM, and the area of the nascent ECM was normalized to the cell area. Aspect ratio, which is the ratio of the cell’s major axis and minor axis, was quantified from the binarized maximum-intensity z-projections. Cell volume was computed by binarizing the CellBrite channel of the stack with Otsu’s thresholding, calculating the cumulative number of voxels in the stack, and then converting from voxels to μm^3^ using the slice thickness.

### 2.12 Fluorescence Microscopy of Vascular Networks

Rhodamine phalloidin staining was used to quantify vessel formation and DAPI was used to generate a nuclei count at day 7. Cells were encapsulated as described in Section 2.8. After seven days in culture, cells were fixed for one hour at room temperature with 4 v% paraformaldehyde and washed 2x5 min at room temperature with PBS. Following washing, the cells were permeabilized for 2 hours at room temperature with a solution comprised of 0.2 v% Triton X-100 in 2 v% paraformaldehyde, washed for 5 min at room temperature with PBS, and then blocked with a blocking buffer (5 v% BSA) in PBS for 2 hours at room temperature. The blocking buffer was replaced with a staining solution comprised of 1 v% BSA with 2 drops/mL of ActinRed™ 555 ReadyProbes™ Reagent (Rhodamine phalloidin) (Invitrogen) and 1:500 DAPI (Sigma Aldrich), and the staining solution was left on the hydrogels for 48 hours at 4°C. Next, the staining solution was discarded and the hydrogels were washed at room temperature with PBS 3x10 minutes. Samples were placed in #1.5 glass bottom mini petri dishes (Cellvis) and imaged on Nikon AXR-NSPARC with a 20X air objective. Three separate encapsulations were performed. For each condition, three samples were imaged per encapsulation, each sample was imaged in the center of the hydrogel at three different locations, and the z-stacks were each 120 µm thick.

To quantify the number of nuclei in each image taken in section 2.12, ImageJ was used to generate a maximum intensity projection of the DAPI channel, Otsu’s thresholding was used to binarize the image, and the analyze particles command was used to count the nuclei. To analyze the vascular networks, we used a computational pipeline that has been previously developed.^46,47^ In brief, images were converted to 8-bit, had their contrast adjusted, were binarized with Li’s thresholding, and despecked. Next, the binarized images were skeletonized with MATLAB, converted into a network comprised of nodes and branches, and analyzed to quantify parameters like volume fraction, vessel diameter, and the total number of branch points.

### 2.13 Gelatin Zymography

HUVECs were encapsulated as described in Section 2.8, and their conditioned media was collected on day 3. To pellet any cellular debris, media samples were centrifuged at 1000 RCF for 10 minutes. Following centrifugation, 500 µL of media supernatant was transferred to the top chamber of a Nanosep centrifugal filter with a 10 K Omega membrane. The samples were centrifuged at 14,000 RCF for 15 minutes at 4°C and then reconstituted with 25 µL of molecular biology grade water. Then, 15 µL of the reconstituted sample was mixed with 15 µL of tris-glycine SDS sample buffer (Invitrogen), and 20 µL of this mixture was loaded into a precast Novex 10% zymogram plus gelatin gel (Invitrogen). The gel was run in an electrophoresis apparatus for 20 minutes at 60 V and then 1.5 hours at 125 V. The gel was then imaged to record the position of the Novex protein standard (Invitrogen) bands. The gel was incubated for thirty minutes with Novex Zymogram Renaturing Buffer (1X, Invitrogen) at room temperature under gentle agitation. The renaturing buffer was replaced with Novex Zymogram Developing Buffer (1X, Invitrogen) and the gel was incubated for thirty minutes at room temperature. Next, the developing buffer was replaced with fresh solution and the gel was incubated overnight at 37°C. The following day, the gel was washed with deionized water 3x5 minutes and then incubated at room temperature overnight under gentle agitation with SimplyBlue SafeStain (Invitrogen). Finally, the gel was imaged to record the position of the bands. The MMP-2 and MMP-9 bands were identified by comparing them to the protein standard and standards for MMP-2 and MMP-9 (Bio Techne). Bands were quantified in accordance with previously published methods.^48^ In summary, the gel image was opened in ImageJ, the rectangular selection tool was used to select lanes, profile plots were generated for each band, and the area under each curve on the profile plot was measured. The area of each curve was normalized to an acellular media control. Media samples were collected from at least three individual hydrogels during two separate encapsulations.

### 2.14 Statistical Analysis

Data comparisons were performed in Prism 10 (GraphPad Software, LLC). Data is reported as mean ± standard deviation. Statistical analysis was performed using a one-way ANOVA followed by a Turkey’s multiple comparisons test. Significance is denoted as the following: *p≤0.05, **p≤0.01, ***p≤0.001, and ****p≤0.0001.

## 3. Results and Discussion

### 3.1 Peptoid substitutions tune hydrogel degradability without altering mechanics

To probe the effects of degradability on HUVEC morphology and vessel formation, we formed NorHA hydrogels (**Fig. 1A**) using a pendant adhesive motif (GCGYGRGDSPG) and library of degradable crosslinkers. The crosslinker library (**Fig. 1B**) ^25,26^consisted of five sequences: a degradable peptide sequence (PAALVA, “Parent”), two sequences with single peptoid substitutions to the Parent sequence in either the P1 (“P1”) or P3 (“P3”) position, a sequence with two peptoid substitutions to the Parent sequence (“Tandem”), and a non-degradable peptoid control (“Peptoid”). The Peptoid crosslinker substituted all the amino acids of the Parent sequence for N-methylglycine peptoid residues. All crosslinkers contained cysteines at the termini to enable thiol-ene crosslinking. Additionally, all crosslinkers contained N-methylglycine spacers to achieve similar sequence lengths to previously established peptide crosslinkers.^18,49^ For all hydrogel conditions, the relative amount of crosslinker was held constant at 25% of the molar amount of norbornenes on the hyaluronic acid.

**Fig. 1.**
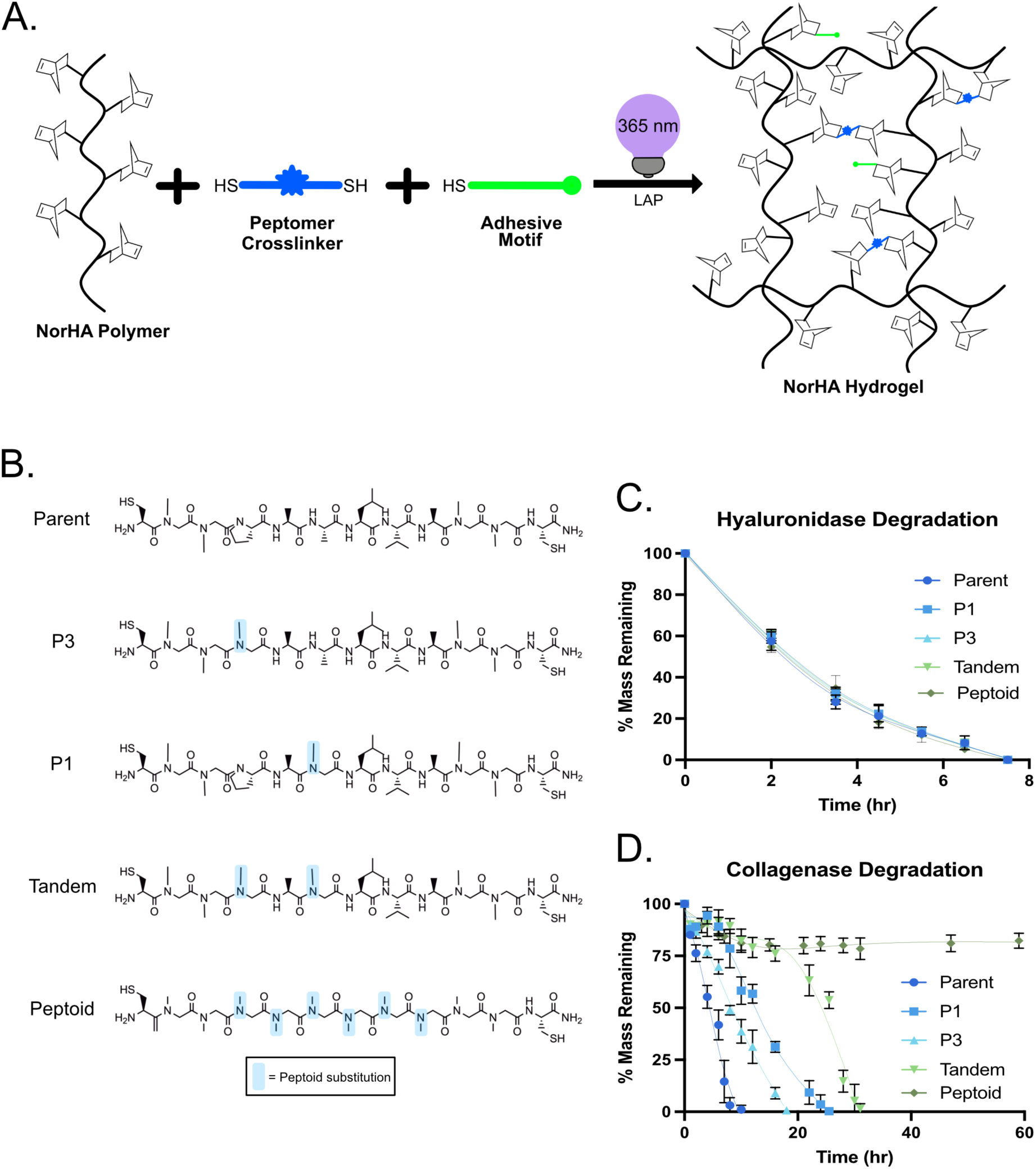
Peptoid substitutions in a peptide crosslinker decreases NorHA hydrogel degradability to collagenase but result in similar hyaluronidase degradation profiles. (A) Schematic of NorHA hydrogel formation during a thiol-ene photopolymerization with LAP and 365 nm light. (B) Peptide and peptide-peptoid hybrid crosslinker structures where the light blue overlay represents a peptoid substitution. (C) Mass retention profiles during hyaluronidase degradation (75 units/mL) over 8 hours. (D) Mass retention profiles during collagenase degradation (250 units/mL) over 60 hours.

We assessed the degradability of our hydrogel formulations by incubating the hydrogels with two different enzyme solutions, hyaluronidase (**Fig. 1C**) and collagenase (**Fig. 1D**), and measuring the mass loss over time. Hyaluronidase degrades the hyaluronic acid backbone of the NorHA hydrogels by cleaving a glycosidic bond.^50^ We observed that all hydrogel formulations displayed similar degradation profiles in response to hyaluronidase and had fully degraded within eight hours. In contrast, collagenase models several MMPs and has been shown to differentially degrade the peptide and peptide-peptoid hybrid sequences used here.^28^ We observed that Parent-crosslinked hydrogels degraded the fastest (within 10 hours), the P1 and P3-crosslinked hydrogels degraded at intermediate rates (within 26 hours and 18 hours respectively), and the Tandem-crosslinked hydrogels degraded the slowest (within 31 hours). The non-degradable Peptoid-crosslinked hydrogels lost approximately 20% of their total mass within the first 10 hours, likely through the loss of unreacted NorHA chains, and then did not degrade further, remaining at a near constant mass for another 50 hours. The trends observed for bulk hydrogel degradation with collagenase align well with published trends for the release rates of similar pendant substrates from a hydrogel with MMP-2 and MMP-9.^28^ Overall, these results suggest that the substitution of peptoids in a peptide crosslinker sequence is a reliable method for decreasing proteolytic susceptibility of hydrogels while maintaining crosslinkers of similar length and structure.

To confirm the various crosslinkers resulted in hydrogels with similar material properties, we used shear oscillatory rheology to measure the hydrogel shear storage modulus, G’ (**Fig. 2A**). All hydrogels were designed to have a G’ around 100 Pa, which has previously been shown to promote endothelial cell spreading and organization.^51–54^ We determined that all the formulations had similar initial storage moduli of approximately 130 Pa. The swelling ratios (**Fig. 2B**) also remained consistent at approximately 60. Thus, this hydrogel platform successfully decoupled degradability from mechanics, adhesive properties, and swelling, rendering it suitable to probe the influence of degradability on *in vitro* vessel formation.

**Fig. 2.**
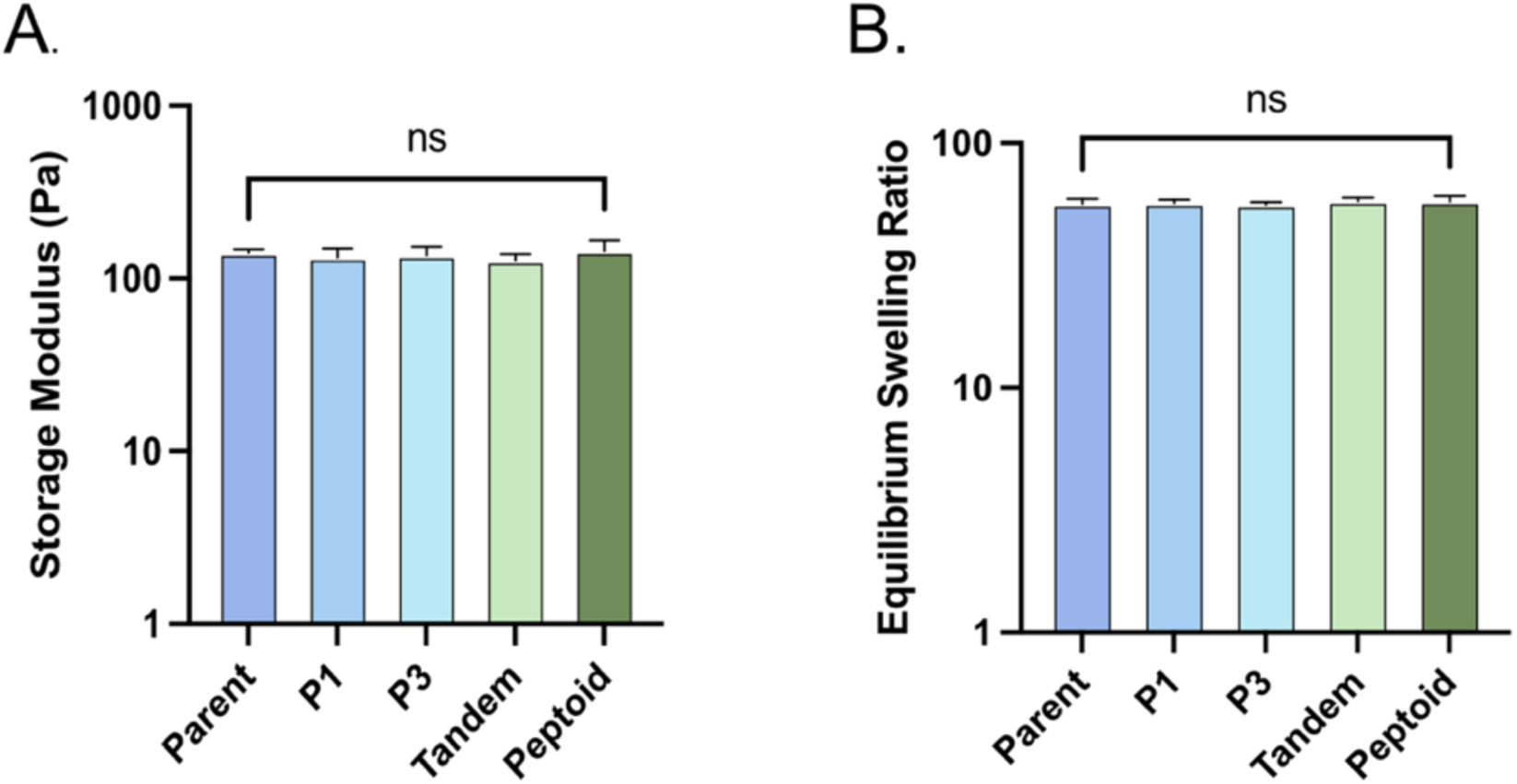
Storage moduli and equilibrium swelling ratios are similar for NorHA hydrogels fabricated with all crosslinkers. (A) Rheological characterization (storage modulus, G’) of NorHA hydrogels crosslinked with the peptide, hybrid, and peptoid crosslinkers. (B) Equilibrium swelling ratio (Qm) of NorHA hydrogels crosslinked with the peptide, hybrid, and peptoid crosslinkers.

### 3.2 Impact of peptoid substitutions on encapsulated HUVEC viability and proliferation

Next, we encapsulated human umbilical vein endothelial cells (HUVECs) at a density of 2.2x10^6^ cells/mL in our hydrogels and evaluated cell viability at several timepoints. One-hour post-encapsulation, we quantified cell viability with a Live/Dead assay and observed over 90% viability for all conditions (**Fig. 3A** and **Fig. 3B**). Additionally, we utilized an MTS assay to assess the metabolic activity of the encapsulated HUVECs at day 1, day 3, and day 7. At day 1, HUVECs encapsulated in all hydrogel formulations had similar levels of metabolic activity (**Fig. 3C**), which suggests that any unincorporated crosslinker did not have an impact on cell viability. By day 3, differences in metabolic activity were apparent (**Fig. 3D**). HUVECs encapsulated in the Parent crosslinked hydrogels had the highest metabolic activity (0.78 ± 0.04 a.u.); HUVECs encapsulated in P1, P3, and Tandem crosslinked hydrogels had intermediate metabolic activity (0.57 ± 0.03, 0.56 ± 0.02 a.u., and 0.53 ± 0.02 a.u., respectively); and the HUVECs encapsulated in Peptoid-crosslinked hydrogels had the lowest metabolic activity (0.47 ± 0.02 a.u.). These differences in metabolic activity were further exacerbated by day 7 (**Fig. 3E**), when the HUVECs in the Parent-crosslinked hydrogels increased their metabolic activity (1.61 ± 0.07 a.u.), but those in the Peptoid-crosslinked hydrogels did not (0.46 ± 0.08 a.u.). HUVECs in the P1 and P3 crosslinked hydrogels also increased their metabolic activity compared to day 3 (1.41 ± 0.05 a.u. and 1.38 ± 0.12 a.u., respectively), whereas the Tandem-crosslinked hydrogels showed reduced metabolic activity (0.89 ± 0.17 a.u.). These results suggest that there is a positive correlation between HUVEC metabolic activity and hydrogel degradability in 3D and corresponds with published data for other cell types.^18,55–58^

**Fig. 3.**
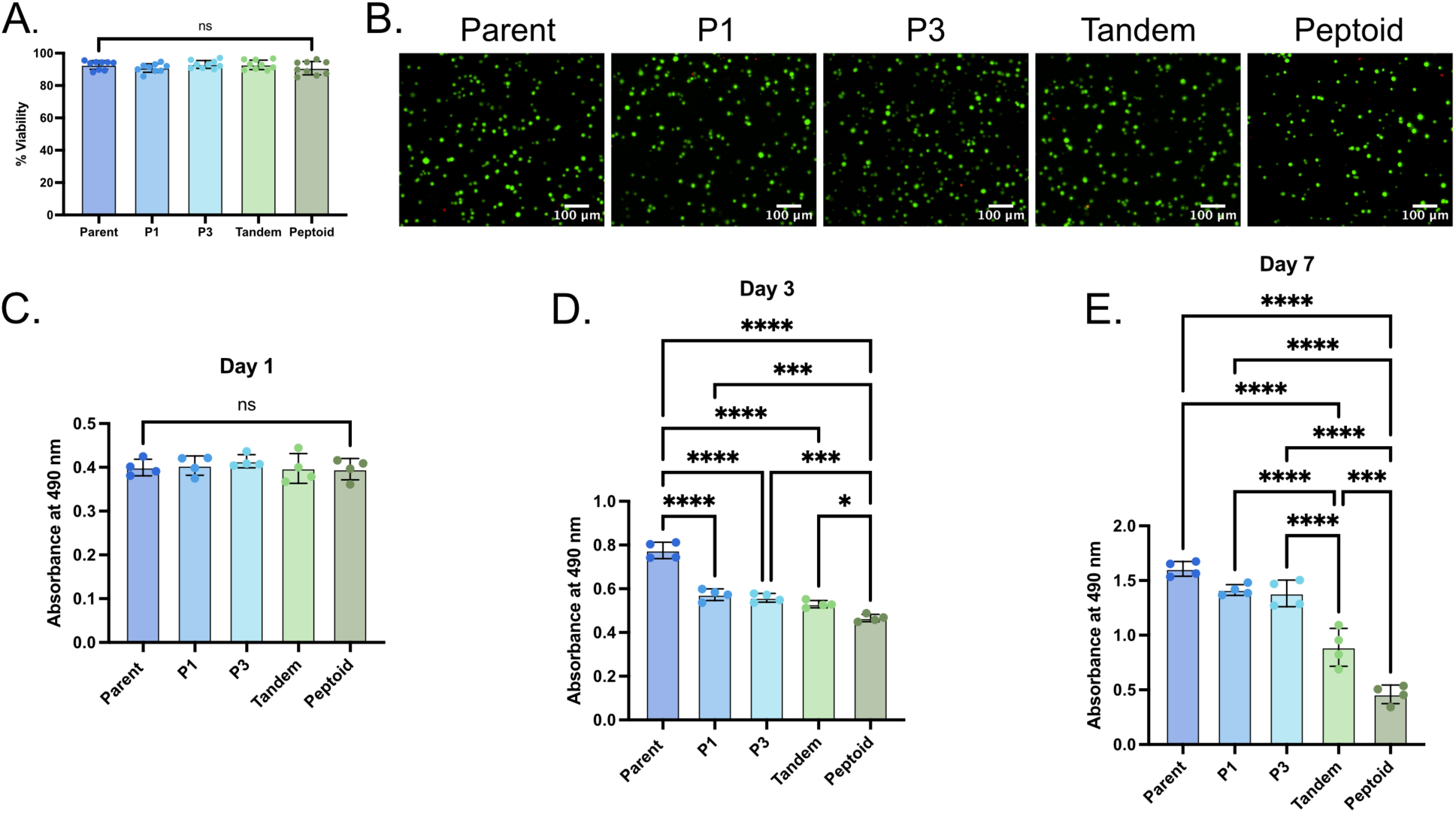
Hydrogel formation with all crosslinkers is cytocompatible, but they result in different amounts of HUVEC proliferation during longer term culture: (A) Quantification of Live/Dead staining shows high viability for all conditions one-hour post-encapsulation, (B) Representative maximum intensity projections of Live/Dead staining of HUVECs 1 hour post encapsulation. Live HUVECs appear green due to calcein-AM cleavage, and dead HUVECs appear red due to membrane permeation of ethidium homodimer-1. Scale bar = 100 μm, (C) Measurement of absorbance at 490 nm for an MTS assay performed on encapsulated HUVECs at day 1, (D) Measurement of absorbance at 490 nm for an MTS assay performed on encapsulated HUVECs at day 3, (E) Measurement of absorbance at 490 nm for an MTS assay performed on encapsulated HUVECs at day 7. Significance is denoted as: *p≤0.05, **p≤0.01, ***p≤0.001, and ****p≤0.0001.

### 3.3 Quantifying HUVEC vessel formation in peptide/peptoid-crosslinked hydrogels

To assess the impact of hydrogel degradability on HUVEC vessel formation, cells were fixed and stained after 7 days in culture (**Fig. 4A** and **Fig. S8**). In line with the metabolic activity results above, we noticed significantly different numbers of cells in the various hydrogels; specifically, the number of cells increased with hydrogel degradability (**Fig. 4B**). There were the most nuclei per image for HUVECs encapsulated in the Parent-crosslinked hydrogels (220 ± 48), intermediate nuclei per image for the P1 and P3-crosslinked hydrogels (149 ± 40 and 125 ± 42, respectively), less nuclei per image for the Tandem-crosslinked hydrogels (70 ± 32), and the least nuclei per image for the Peptoid-crosslinked hydrogels (46 ± 11).

**Fig. 4.**
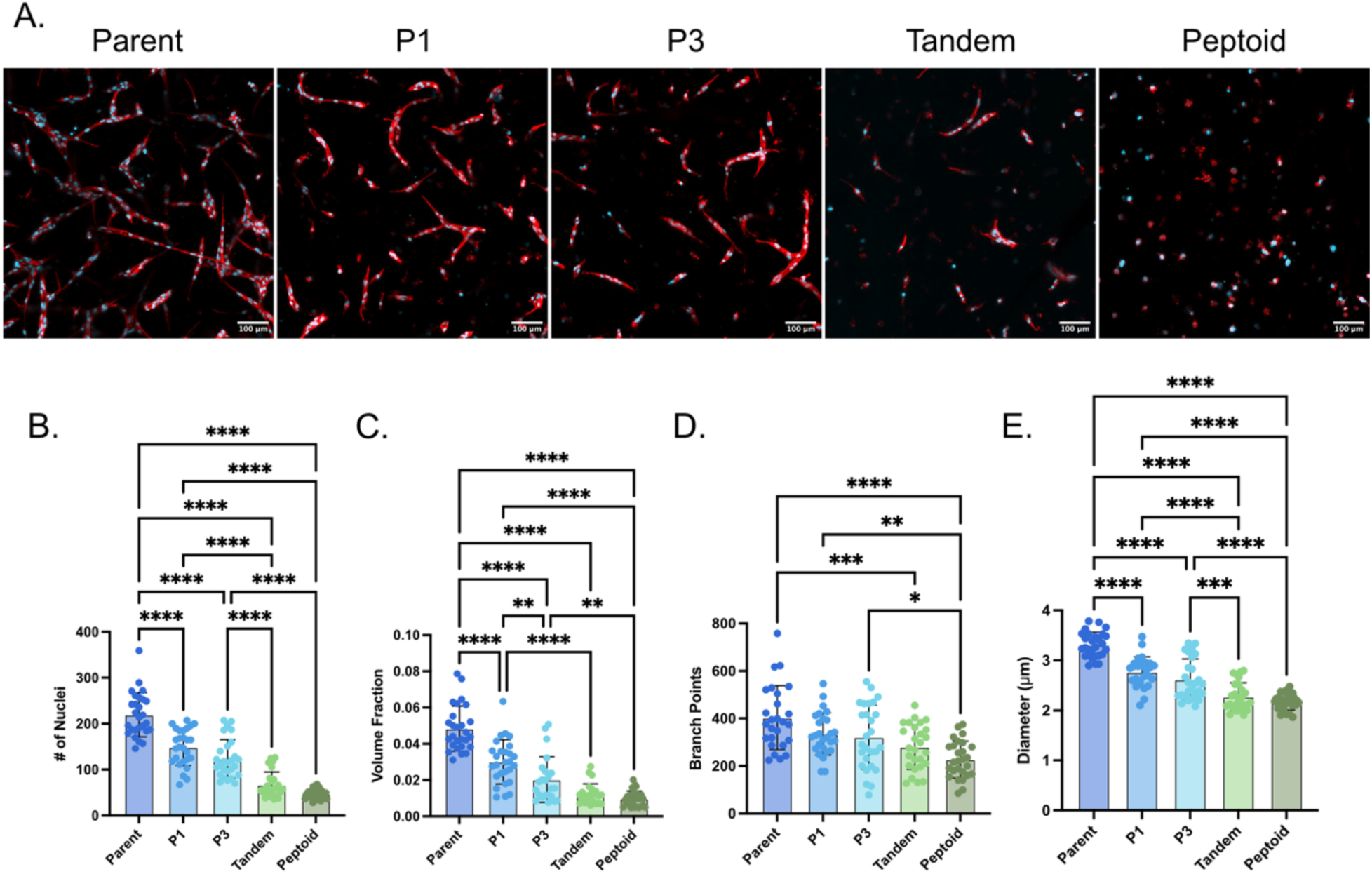
At day 7, HUVECs encapsulated in the most degradable NorHA hydrogels have proliferated the most and have formed the most vessels. (A) Representative maximum intensity projections (Z = 120 µm) of encapsulated HUVECs at day 7 stained for f-actin (red) and nuclei (blue). Scale bar = 100 µm. (B) Number of nuclei at day 7, (C) Vessel volume fraction at day 7, (D) Number of branch points at day 7, (E) Vessel diameter in µm. Significance is denoted as: *p≤0.05, **p≤0.01, ***p≤0.001, and ****p≤0.0001.

In addition, we observed similar trends with vessel formation. After a week of culture, HUVECs encapsulated in more degradable hydrogels formed more vessel-like structures than those encapsulated in less degradable hydrogels or non-degradable hydrogels (**Fig. 4A**). Quantification via a computational pipeline confirmed this result by calculating the volume fraction (which is the fraction of the hydrogel that is occupied by the vascular network; **Fig. 4C**), number of branch points (**Fig. 4D**), and average vessel diameter (**Fig. 4E**).^46,47^ HUVECs encapsulated in Parent-crosslinked hydrogels had the greatest volume fraction (0.048 ± 0.012), most branch points (403 ± 134), and largest average diameter (3.31 ± 0.25 µm). In the less degradable P1 and P3 crosslinked hydrogels, HUVECs had a lower volume fraction (0.030 ± 0.012 and 0.020 ± 0.013, respectively), fewer branch points (335 ± 89 and 321 ± 135, respectively), and a smaller average diameter (2.77 ± 0.30 µm and 2.62 ± 0.41 µm, respectively). In the least degradable Tandem-crosslinked hydrogels, HUVECs exhibited further reduction in volume fraction (0.012 ± 0.006), the number of branch points (280 ± 95), and average diameter (2.28 ± 0.28 µm). Finally, HUVECs encapsulated in the non-degradable, Peptoid-crosslinked hydrogels did not form any visible vessel-like structures. We hypothesize that the decrease in vessel formation observed in the less-degradable hydrogels is due to the reduced metabolic activity and lower cell number quantified via MTS assay and nuclei count. This result highlights the importance of protease-mediated matrix remodeling in the vascularization of tissue constructs.^15,17,59,60^

### 3.5 MMP-2 and MMP-9 secretion from HUVECs encapsulated in peptide, peptomer, and peptoid crosslinked hydrogels

During vasculogenesis, proteolytic matrix degradation occurs via cell secretion of MMPs, particularly MMP-2 and MMP-9, which allows for cell spreading and for the release of ECM bound cytokines that are directly involved in vasculogenesis.^7,38,39^ Thus, we assessed the effect of hydrogel degradability on the proteolytic activity of MMP-2 and MMP-9 via gelatin zymography. Conditioned media from all hydrogel conditions revealed degradation of the parent substrate by MMP-2 and MMP-9, with more intense bands corresponding to conditioned media from less degradable hydrogels (**Fig. 5A** and **Fig. S9**). Quantification of the bands confirmed this result; conditioned media from the least degradable Tandem-crosslinked hydrogels and the non-degradable Peptoid-crosslinked hydrogels had fold-change increases over the Parent-crosslinked hydrogels of approximately 1.3 for expression of pro-MMP-9 (**Fig. 5B**), 1.4 for expression of active MMP-9 (**Fig. 5C**), 1.5 for the expression of pro-MMP-2 (**Fig. 5D**), and 1.3 and 2.0, respectively, for expression of active MMP-2 (**Fig. 5E**). This difference is further exacerbated when considering the amount of active MMP-2 and MMP-9 on a per cell basis. At the time that the conditioned media was collected there are significantly more metabolically active HUVECs in the Parent-crosslinked hydrogels than the Tandem-crosslinked hydrogels and the Peptoid-crosslinked hydrogels (**Fig. 3D**), so the amount of MMP-2 and MMP-9 secreted on a per cell basis in the Parent-crosslinked hydrogels is drastically lower than in the Tandem-crosslinked hydrogels and the Peptoid-crosslinked hydrogels.

**Fig. 5.**
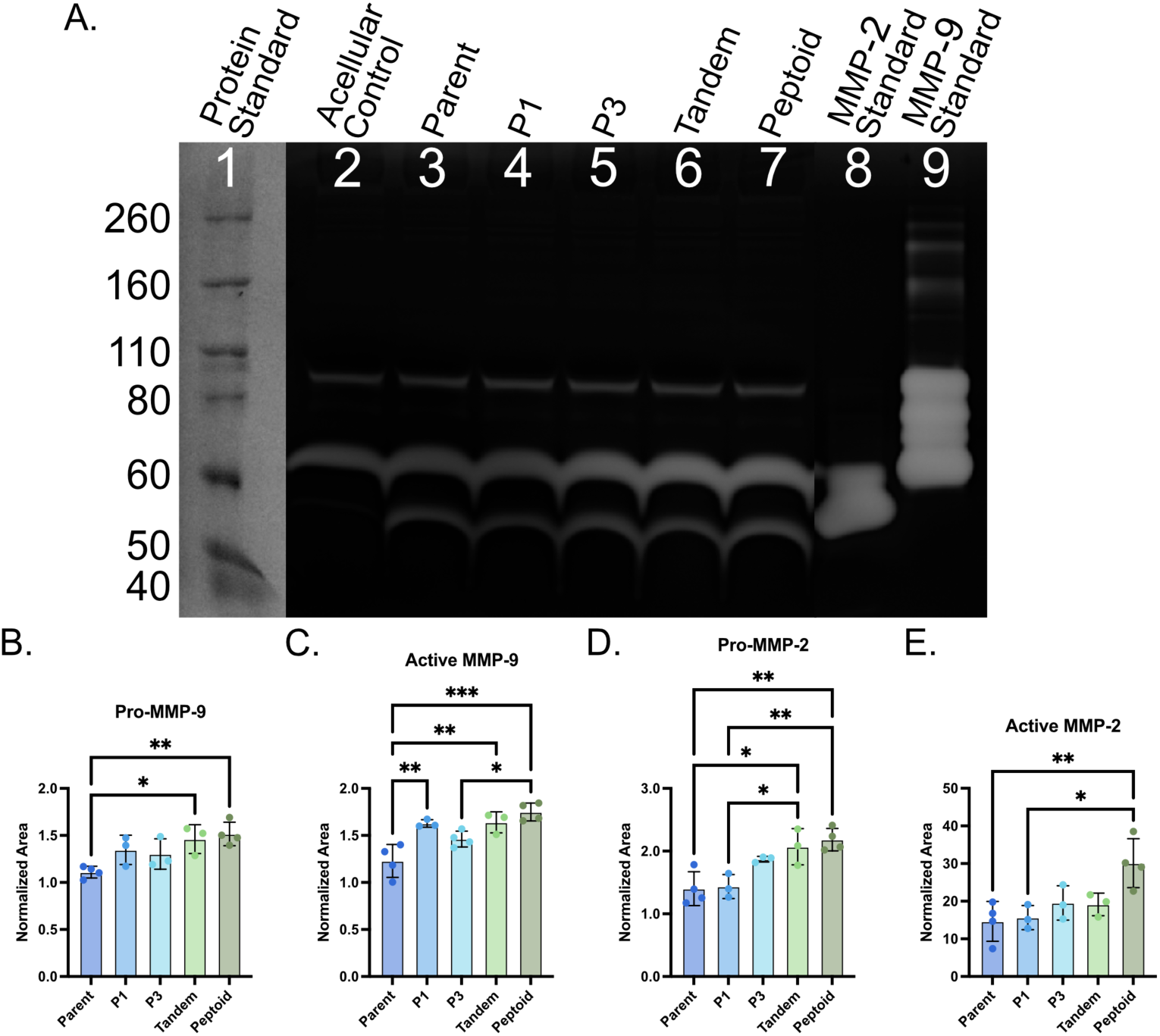
Based on gelatin zymography, HUVECs encapsulated in less degradable NorHA hydrogels secrete more MMP-9 and MMP-2 on day 3. (A) Scanned image of gelatin zymogram where the lane numbers correspond to the following: 1 is a protein standard, 2 is an acellular media control, 3 is conditioned media from a parent hydrogel, 4 is conditioned media from a P1 hydrogel, 5 is conditioned media from a P3 hydrogel, 6 is conditioned media from a tandem hydrogel, 7 is conditioned media from a peptoid hydrogel, 8 is a MMP-2 standard, and 9 is a MMP-9 standard. (B) Area of Pro-MMP-9 peaks normalized to the area of the acellular media control (C) Area of Active MMP-9 peaks normalized to the area of the acellular media control (D) Area of Pro-MMP-2 peaks normalized to the area of the acellular media control (E) Area of Active-MMP-2 peaks normalized to the area of the acellular media control. Significance is denoted as: *p≤0.05, **p≤0.01, ***p≤0.001, and ****p≤0.0001.

This result suggests that the HUVECs encapsulated in less degradable hydrogels increased their MMP expression in an attempt to degrade the surrounding matrix. However, the reduced proteolytic susceptibility of the Tandem-crosslinked hydrogels and the lack of proteolytic susceptibility of the Peptoid-crosslinked hydrogels prevented protease-mediated remodeling of HUVECs into vessel-like structures (**Fig. 4**). These results are corroborated by previous work that evaluated the MMP expression and vasculogenesis of another type of endothelial cell (endothelial colony-forming cells (ECFCs)) in hydrogels that were only crosslinked with MMP-degradable peptides and hydrogels that were crosslinked with a combination of MMP-degradable peptides and non-degradable kinetic chains.^61^ The ECFCs encapsulated in the hydrogels that contained non-degradable kinetic chains expressed higher levels of MMP but had inhibited vascular formation in comparison to the hydrogels containing only MMP-degradable crosslinks.^61^ Collectively, these data suggest that although MMP-mediated remodeling is important for vasculogenesis, higher expression of MMP-2 and MMP-9 is not necessarily correlated with enhanced vessel formation and could be indicative of an environment that limits protease-mediated degradation.

### 3.6 Morphology and ECM secretions for HUVECs encapsulated in peptide/peptoid-crosslinked hydrogels

During vessel formation, individual endothelial cells assemble and, along with supporting cells, deposit a form of ECM called the basement membrane, which is comprised primarily of proteins (like collagen) and glycoproteins (like laminin and fibronectin).^3,6,40,62^ This basement membrane provides organizational cues to endothelial cells, structural support to vessels, and serves as a reservoir for growth factors that directly influence vasculogenesis.^6,40^ Hence, we explored the effect of hydrogel degradability on HUVEC single cell morphology and the deposition of nascent ECM. To do this, we encapsulated HUVECs in our differentially degradable hydrogels at a density of 1x10^5^ cells/mL and employed a bio-orthogonal technique for the metabolic labelling of glycans, which fluorescently labels nascent glycoproteins post-synthesis and secretion.^42,43^

On day 3, HUVECs encapsulated in the Parent-crosslinked hydrogels appeared larger and more spread than HUVECs encapsulated in P1 and P3-crosslinked hydrogels, but all exhibited a spindle-like morphology (**Fig. 6A**). However, HUVECs encapsulated in the Tandem and Peptoid-crosslinked hydrogels appeared smaller and mostly rounded. Quantification of volume (**Fig. 6B**) and aspect ratio (**Fig. 6C**), which is a metric of cell spreading, aligned well with this observation. HUVECs encapsulated in the Parent-crosslinked hydrogels had the largest average volume (5,500 ± 2,000 μm^3^); HUVECs cultured in the P1 and P3-crosslinked hydrogels had intermediate volumes (4,800 ± 2,000 μm^3^ and 4,600 ± 1,600 μm^3^, respectively); and HUVECs encapsulated in the Tandem and Peptoid-crosslinked hydrogels had the smallest average volumes (3,600 ± 1,200 μm^3^ and 2,700 ± 700 μm^3^, respectively). Similarly, the average aspect ratio for HUVECs encapsulated in the Parent-crosslinked hydrogels was the largest (4.11 ± 1.91); the average aspect ratio for the P1 and P3-crosslinked hydrogels were intermediate (2.53 ± 1.25 and 2.56 ± 1.15, respectively); and the average aspect ratio of HUVECs encapsulated in the Tandem and Peptoid-crosslinked hydrogels were the smallest (1.46 ± 0.34 and 1.33 ± 0.27, respectively).

**Fig. 6.**
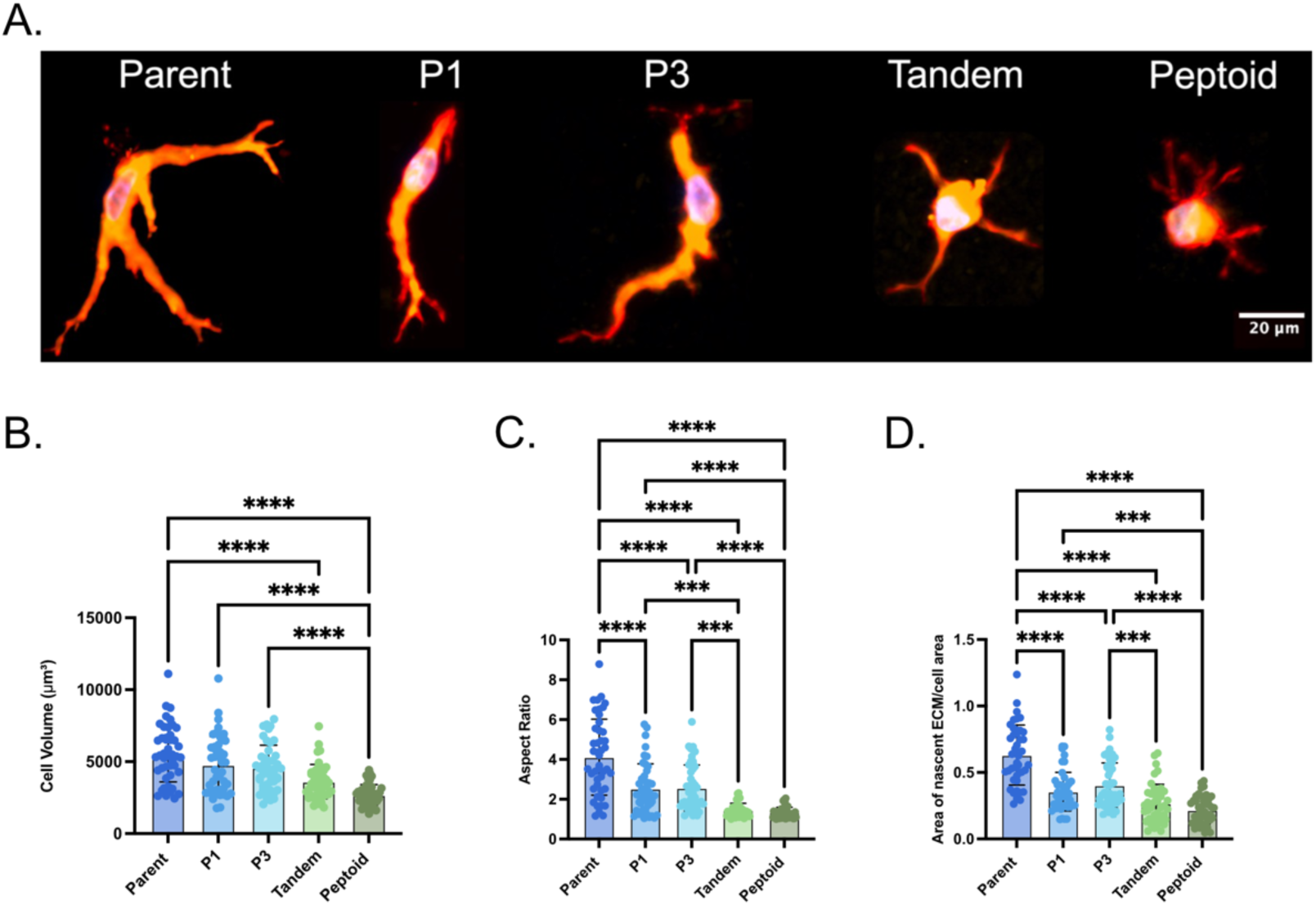
HUVECs encapsulated in more degradable NorHA hydrogels have the largest volume, spread the most, and secrete the most nascent ECM. (A) Representative maximum intensity projections of encapsulated HUVECs at day 3 stained for nascent ECM secretions (red), cell membrane (yellow), and nuclei (blue). Scale bar = 20 μm. (B) HUVEC volume at day 3 in μm^3^, (C) HUVEC aspect ratio at day 3, (D) Area of ECM secretion thickness normalized to cell area at day 3. Significance is denoted as: *p≤0.05, **p≤0.01, ***p≤0.001, and ****p≤0.0001.

After analyzing cell morphology, we quantified the amount of nascent ECM secretions and normalized them to the cell area to account for differences in cell size; this normalization resulted in a ratio of the area of ECM secretions to cell area. The amount of ECM secretions followed a similar trend to morphology metrics, and we observed more ECM deposition in more degradable hydrogels. The HUVECs encapsulated in Parent-crosslinked hydrogels had significantly more ECM deposition than all other conditions (0.63 ± 0.23), the HUVECs encapsulated in P1 and P3-crosslinked hydrogels had intermediate ECM deposition (0.36 ± 0.15 and 0.40 ± 0.17, respectively), and the HUVECs encapsulated in the Tandem and Peptoid-crosslinked hydrogels had the least ECM deposition (0.27 ± 0.14 and 0.22 ± 0.10, respectively). Overall, these results suggest that reducing proteolytic susceptibility of the hydrogels limits the ability of the encapsulated HUVECs to remodel into larger, more spread morphologies and restricts the secretion of nascent ECM. Furthermore, this decrease in cell spreading and lack of ECM deposition due to lessened hydrogel degradability align with the trends observed in vessel formation (**Fig. 4**) where the ability of HUVECs to organize into vessel-like structures was reduced in the less degradable P1, P3, Tandem-crosslinked hydrogels and the non-degradable Peptoid-crosslinked hydrogels.

## 4. Conclusion

Here, we provide a strategy for independently tuning hydrogel proteolytic degradability in vasculogenic scaffolds through the introduction of peptoids in our crosslinking sequences. Our findings highlight the importance of protease-mediated matrix remodeling for vessel formation and show a direct correlation between degradability and vasculogenic potential. Overall, we observed that HUVECs encapsulated in the most degradable (Parent) hydrogels proliferated significantly more, had higher metabolic activity, and formed significantly more vessel-like structures after a week of culture than HUVECs encapsulated in less degradable (P1 and P3) hydrogels. Even fewer vessel-like structures, less proliferation, and less metabolic activity was observed for HUVECs encapsulated in the least degradable (Tandem) hydrogels. HUVECs encapsulated in the non-degradable control (Peptoid) hydrogels show no vessel formation. Single-cell morphology metrics, specifically volume and aspect ratio, and the quantification of nascent ECM secretions aligned well with these trends; cell volume, aspect ratio, and the amount of secreted ECM increased with degradability. Contrastingly, HUVECs encapsulated in more degradable hydrogels expressed less MMP-2 and MMP-9 than HUVECs encapsulated in the less degradable and non-degradable hydrogels, suggesting a feedback loop between the cells and the matrix properties. These data indicate a relationship between hydrogel degradability and the extent to which HUVECs can assemble into vessels and suggest that peptomer crosslinkers might be an effective way to guide vascularization *in vitro* by tailoring scaffold degradability. Furthermore, although the most degradable (Parent) hydrogels facilitated the most vessel formation, the vessel formation is still lacking in regard to volume fraction and connectivity. Future work could further enhance vascularization by incorporating a support cell type (like fibroblasts or mesenchymal stem cells) to increase the secretion of growth factors and ECM components that guide the formation and stabilization of vessels;^15,63–65^ encapsulating nanofibers within the hydrogels to provide mechanical cues that enhance cell spreading and cell-matrix interactions;^66,67^ or utilizing a hydrogel that preserves bioactive ligands and microstructure from the native ECM while still allowing mechanical tunability and stability, like norbornene-modified decellularized ECM.^68^

## 5. Acknowledgments

This research was funded by the National Heart, Lung, Blood Institute of the National Institutes of Health (R01HL157829). This work was also supported by the National Institutes of Health (R35GM138193). MALDI-TOF was performed at the UT Austin Center for Biomedical Research Support Biological Mass Spectrometry Facility (RRID: SCR_021728). Confocal microscopy was performed at the Center for Biomedical Research Support Microscopy and Imaging Facility at UT Austin (RRID: SCR_021756).

