## Supplemental Information for "Tuning Scaffold Degradation with Non-Natural Peptidomimetics to Control Human Umbilical Vein Endothelial Cell Morphology and Vessel Formation"

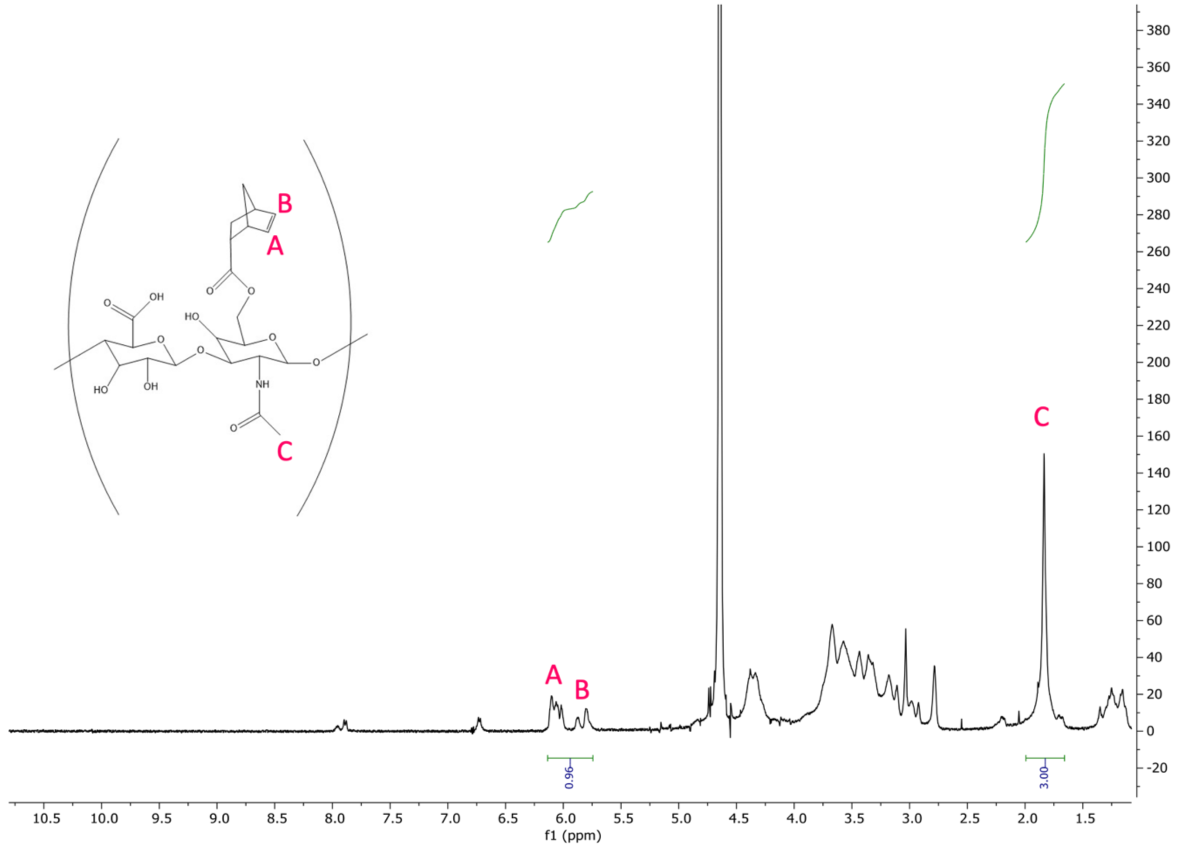


**Figure S1:** $^1$H NMR spectrum of NorHA in deuterium oxide ($D_2O$) demonstrating a degree of functionalization 48%.

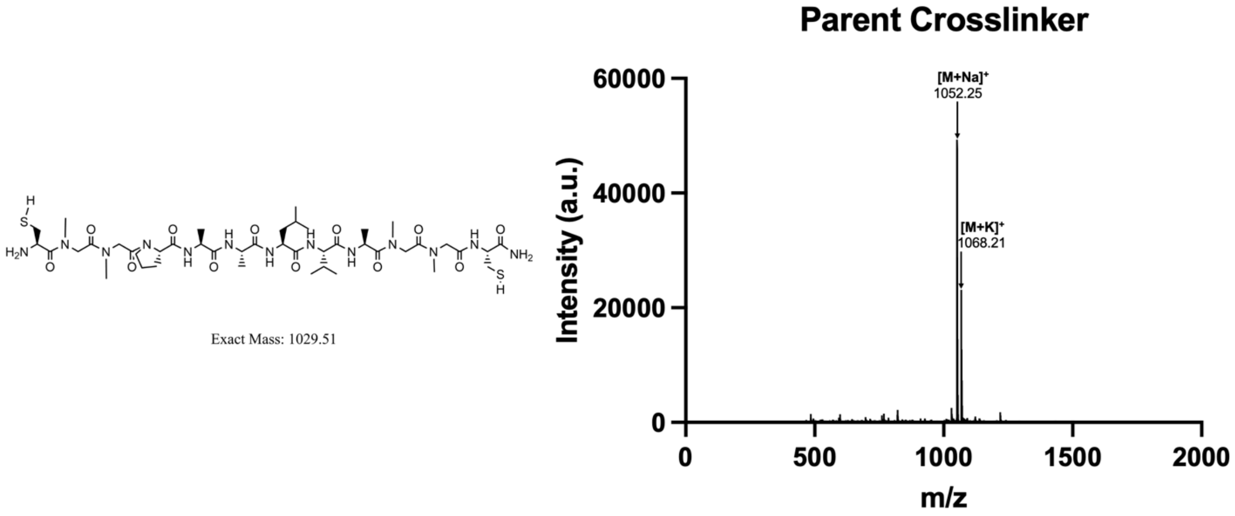


**Figure S2:** Parent crosslinker structure and MALDI data confirming the molecular weight

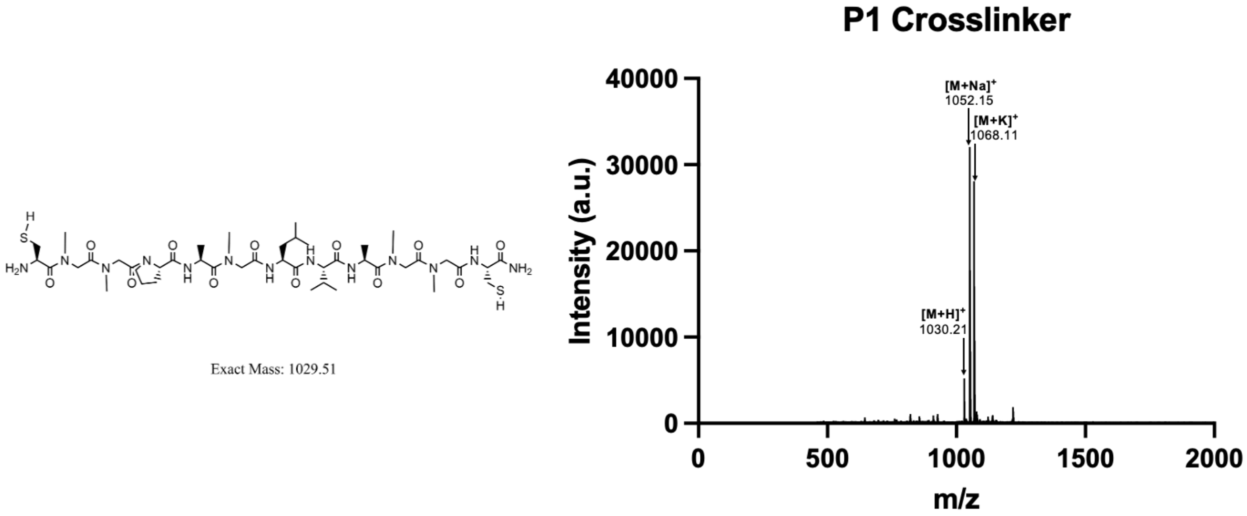


**Figure S3:** P1 crosslinker structure and MALDI data confirming the molecular weight

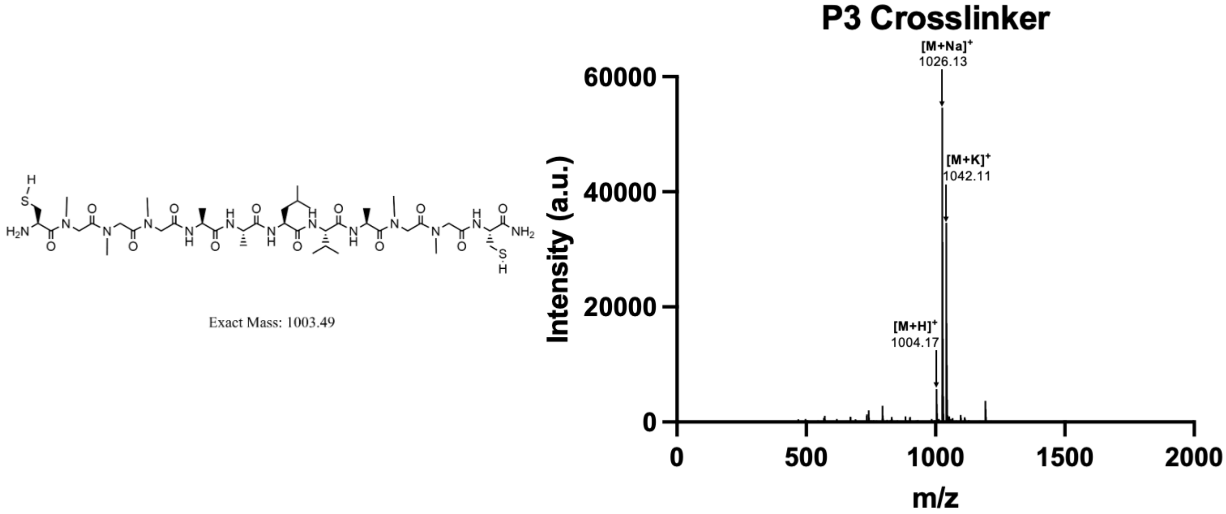


**Figure S4:** P3 crosslinker structure and MALDI data confirming the molecular weight

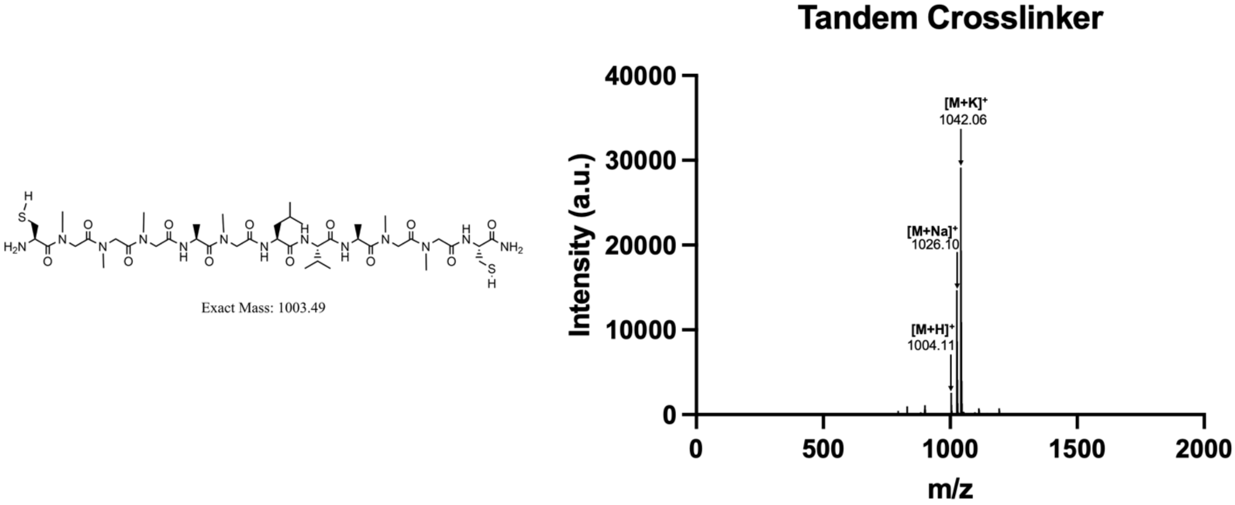


**Figure S5:** Tandem crosslinker structure and MALDI data confirming the molecular weight

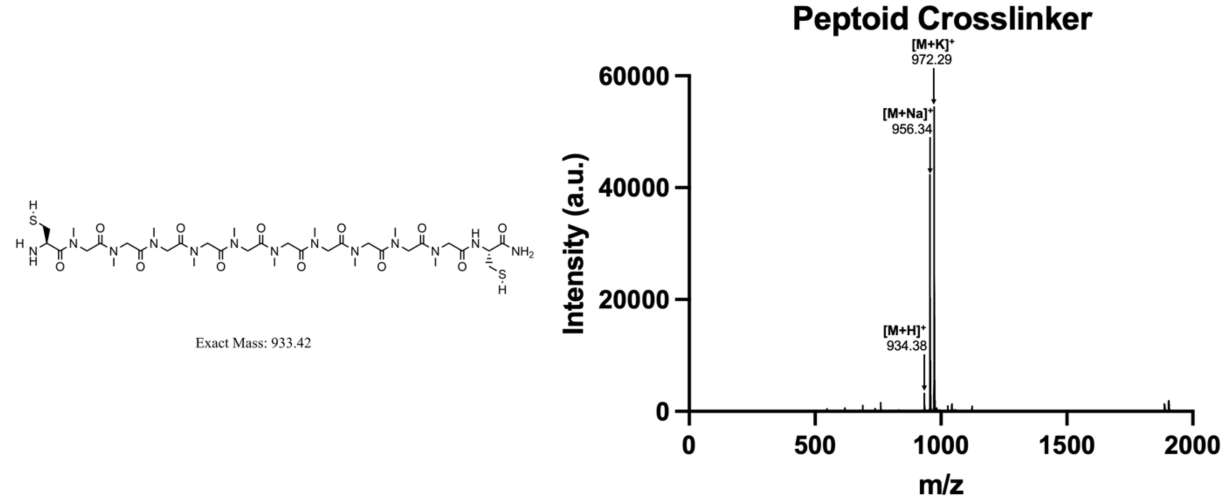


**Figure S6:** Peptoid crosslinker structure and MALDI data confirming the molecular weight

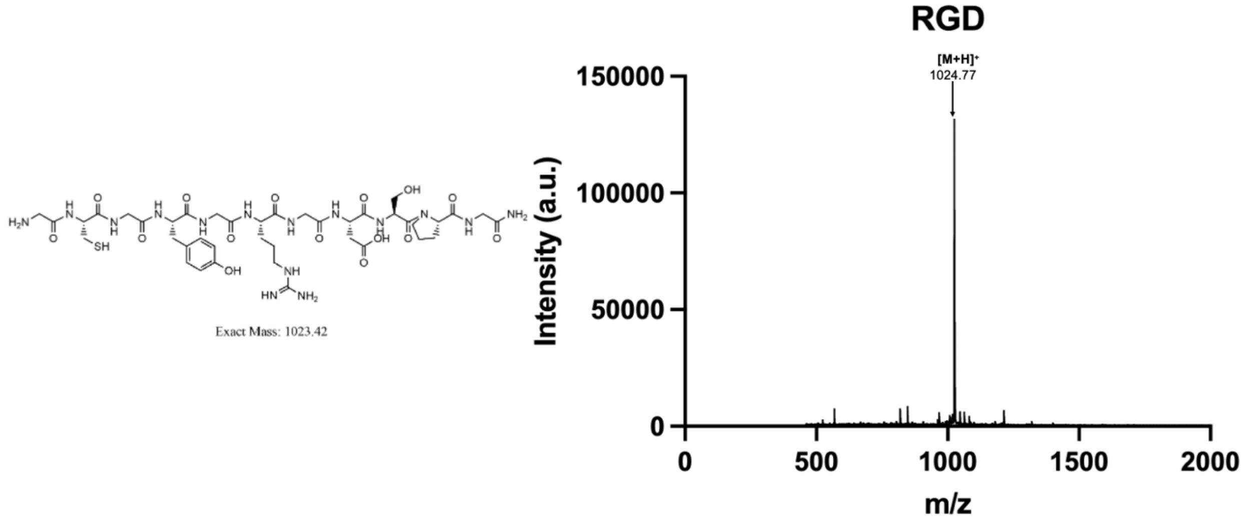


**Figure S7:** RGD cell adhesive motif structure and MALDI data confirming the molecular weight

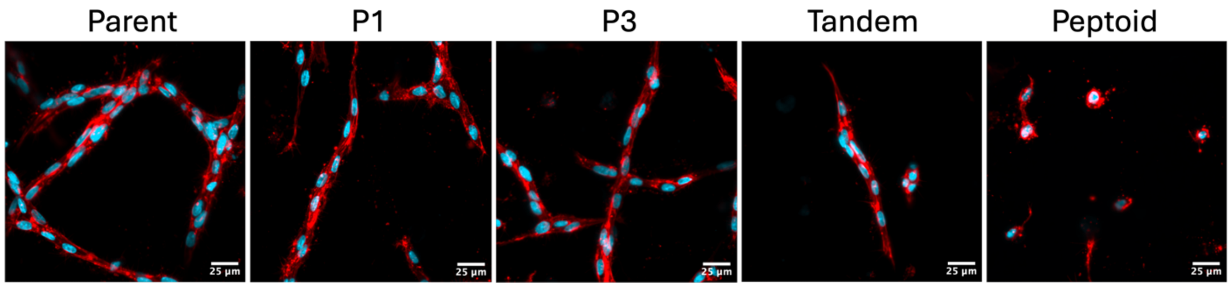


**Figure S8:** Encapsulated HUVECs at day 7 that are stained for f-actin (red) and nuclei (blue). Scale bar = 25 µm.

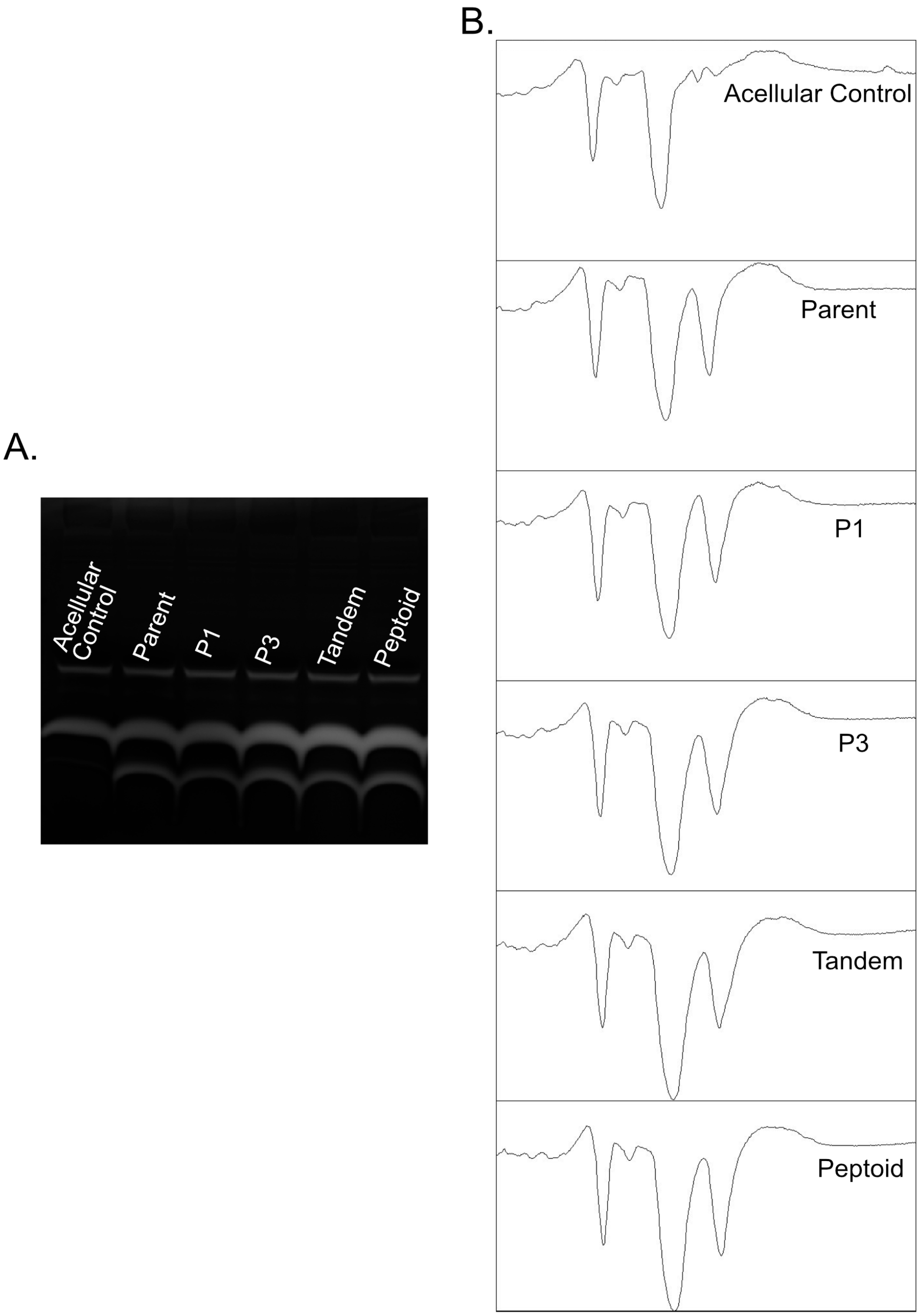


**Figure S9:** (A) Scanned image of gelatin zymogram (B) A profile plot from ImageJ where each panel corresponds to a lane in the scanned zymogram
